# ProMeta: A meta-learning framework for robust disease diagnosis and prediction from plasma proteomics

**DOI:** 10.64898/2026.01.28.702242

**Authors:** Han Li, Haoteng Gu, Lei Hu, Zimo Zhang, Yongji Lv, Peng Gao, Johnathan Cooper-Knock, Yaosen Min, Jianyang Zeng, Sai Zhang

**Affiliations:** School of Mathematical Sciences and LPMC, Nankai University, Tianjin, China; Sheffield Institute for Translational Neuroscience, University of Sheffield, Sheffield, UK; School of Computer Science, Beijing University of Technology, Beijing, China; Department of Artificial Intelligence, School of Engineering, Westlake University, Hangzhou, China; School of Life Sciences, Westlake University, Hangzhou, China; Delta College, Westlake Residential College, Westlake University, Hangzhou, China; Department of Environmental Health, Harvard T.H. Chan School of Public Health, Boston, USA; Zhongguancun Institute of Artificial Intelligence, Beijing, China; Center for Interdisciplinary Studies, School of Science, Westlake University, Hangzhou, China; Department of Biomedical Informatics & Data Science, Yale School of Medicine, New Haven, CT, USA

## Abstract

The plasma proteome offers a dynamic window of human health, capturing the real-time intersections between genetics and physiology. However, the application of deep learning to proteomics is currently hindered by a reliance on large-scale labeled datasets, rendering standard models ineffective for rare or novel diseases where patient samples are inherently scarce. Here, we present ProMeta, a meta-learning framework designed to enable robust disease modeling under extreme data restrictions. By integrating knowledge-guided pathway encoding with bi-level meta-optimization, ProMeta projects unstructured proteomic profiles into biologically interpretable functional tokens. This architecture allows the model to learn a global initialization containing transferable biological priors from biobank-scale data, facilitating rapid adaptation to novel tasks. Through comprehensive benchmark experiments, ProMeta consistently outperformed transfer learning and traditional machine learning baselines in both disease diagnosis and prediction tasks. In the most challenging 4-shot scenarios (utilizing only 2 cases and 2 controls), the model achieved robust generalization with an average AUROC of ∼ 0.69, representing a 24.6% relative improvement over the best-performing baseline methods. Mechanistic investigation revealed that ProMeta disentangles cases from controls in the latent space prior to task-specific adaptation, confirming the acquisition of universal biological rules rather than rote memorization. Furthermore, gradient-based interpretation identified disease-specific protein biomarkers and functional pathways consistent with known pathophysiology. Collectively, ProMeta overcomes the data-scarcity bottleneck in precision medicine, providing a scalable, interpretable framework for characterizing the full spectrum of human diseases, particularly for rare conditions lacking extensive clinical cohorts.

## Introduction

The translation of molecular biology into precision medicine is increasingly dependent on the analysis of the plasma proteome — the comprehensive set of proteins circulating in the blood. Unlike the genome, which offers a static view of disease risk, the proteome is dynamic, capturing the real-time intersections of genetic predisposition, environmental exposure, and pathological progression^[1,2,3,4]^. In recent years, large-scale initiatives such as the UK Biobank (UKB) have generated large-scale high-throughput plasma proteomics profiles for tens of thousands of individuals, providing an unprecedented resource for systematically characterizing proteome–disease associations^[5,6]^.

Proteomic data has been proven powerful to characterize systemic biological aging and organ-specific health. Specifically^[7,8,9,10,11,12]^, Argentieri et al. developed a proteomic aging clock based on nearly 3,000 proteins, showing that accelerated proteomic aging is strongly associated with mortality and the incidence of several chronic diseases across diverse populations^[9]^. Recent studies have also established the utility of plasma proteomics in broad disease contexts. Deng et al. comprehensively mapped 2,920 plasma proteins to 406 prevalent and 660 incident diseases in over 53,000 adults, identifying over 160,000 protein-disease associations^[10]^. Li et al. proposed Prophet, a multistage, multitask transformer-based framework trained on personal proteomes achieving significant gains in the diagnostic and predictive accuracy of common diseases compared to traditional machine learning baselines^[12]^.

However, a critical limitation persists: existing state-of-the-art methods predominantly rely on supervised learning paradigms that require extensive labeled data for every target condition. While successful for common phenotypes with thousands of cases, these approaches struggle to generalize to rare diseases or novel conditions where patient samples are inherently scarce. Consequently, the rapid adaptation of proteomic prediction models to new disease tasks under severely constrained sample conditions remains an unsolved problem, representing a critical bottleneck that limits the extension of proteomics-based predictive frameworks to the “long tail” of human diseases.

To address this challenge, we propose ProMeta, a meta-learning framework specifically designed to leverage the wealth of biobank-scale data to enable robust few-shot diagnosis and prediction for rare and data-scarce conditions. Unlike traditional transfer learning, which optimizes for a specific set of fixed tasks, ProMeta adopts a “learning-to-learn” paradigm. By treating each disease as a distinct task during a bi-level optimization process, the model learns a highly transferable global initialization — effectively encoding universal biological priors into its latent space. This allows ProMeta to rapidly adapt to novel diseases using as few as four patient samples (4-shot learning). We demonstrate that ProMeta significantly outperforms state-of-the-art baselines in both prevalent disease diagnosis and incident risk prediction, particularly in extreme low-data regimes. Furthermore, by integrating gradient-based attribution with biological knowledge graphs, ProMeta offers interpretable insights into the molecular mechanisms driving its predictions, identifying disease-specific biomarkers and pathways that align with known pathophysiology.

## Methods

### 2.1. Data collection and preprocessing

We analyzed plasma proteomics data from the UKB-PPP^[6]^, covering 2,923 proteins across 53,014 UK Biobank^[5]^ participants. Diagnoses were extracted from UK Biobank inpatient records (Fields 41270/41280) using ICD-10 ^[13]^ codes. Following established methods^[10,12]^, we applied FinnGen^[14]^ Data Freeze 12 definitions and quality control guidelines. Cases were classified as prevalent or incident relative to the baseline blood draw. To ensure strict control definitions, we excluded prior diagnoses from incident analyses and future diagnoses (follow-up) from prevalent analyses. Thus, controls were defined as participants free of the disease throughout the study. After removing entries with missing data, the final cohort included 53,014 participants, with normalized expression levels of 2,923 proteins used as model inputs.

### 2.2. Problem formulation

We formulate disease prediction as a few-shot learning framework. Let 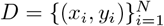 denote the dataset comprising *N* participants, where *x*_*i*_ denotes the proteomic features of the i-th participant and *y*_*i*_ represents the corresponding disease label. Adopting a meta-learning paradigm, our objective diverges from standard supervised learning. Instead of minimizing loss over specific instances, we aim to train a model across a distribution of training diseases *T*_*train*_ to learn transferable meta-knowledge. This enables the model to rapidly adapt to unseen diseases *T*_*test*_ using only limited support examples.

### 2.3. The ProMeta framework

To solve this task, we propose the ProMeta framework (Fig. 1), designed to systematically model the biological heterogeneity underlying proteomic data by encoding unstructured protein expression profiles into a unified pathway representation that captures both molecular specificity and biological interactions. Through bi-level meta-optimization, ProMeta learns comprehensive disease-agnostic meta-knowledge, facilitating rapid adaptation to novel clinical conditions with limited sample availability.

**Fig. 1:**
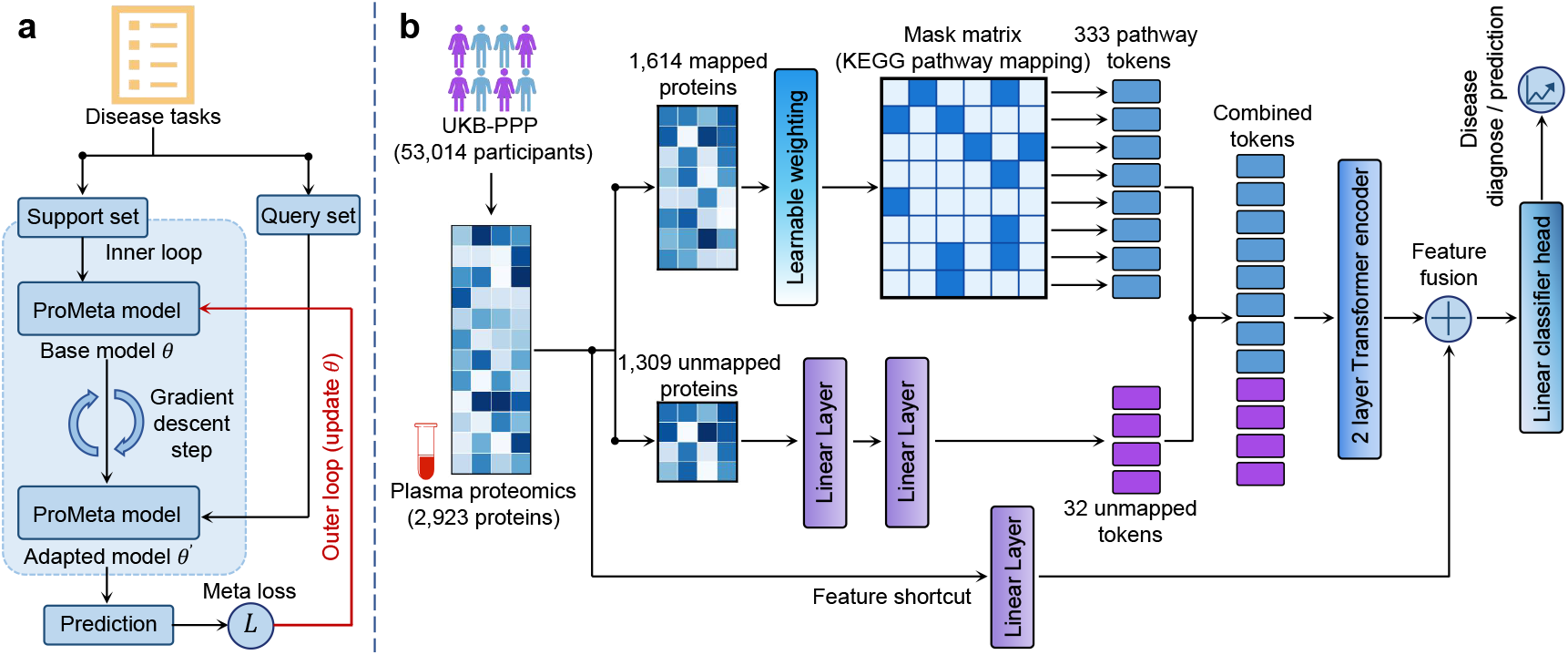
Overall architecture of ProMeta for few-shot disease diagnosis and prediction from plasma proteomics. (a) Meta-learning algorithm. The model employs a bilevel optimization scheme. It uses the support set to adapt to specific disease tasks via gradient descent (inner loop) and leverages the query set to update global parameters (outer loop), ensuring robust generalization to unseen diseases. (b) The network uses a dual-view representation to encode plasma profiles. Proteins are aggregated into pathway-aware tokens using a prior knowledge mask and processed via a Transformer encoder. A residual shortcut connects the original input features directly to the final classification head, combining global pathway context with local protein-level signals.

#### 2.3.2. Hierarchical representation learning

##### 2.3.2.1. Construction of biologically-informed pathway tokens

To accurately model the functional state of biological systems, we encode the input protein expression profile *x ∈ ℝ*^*P*^ (where *P* denotes the number of proteins) into a set of pathway-aware tokens using a knowledge-guided weighted aggregation strategy.

Adaptive feature modulation. First, to mitigate biases from ubiquitous noise and dynamic variation in proteomics data, we project each protein scalar *x*_*j*_ into a high-dimensional latent space and apply an adaptive modulation mechanism. Specifically, for the *j*-th protein, we compute a modulated embedding 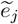 :

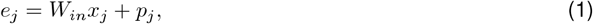

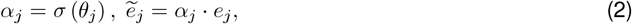

where *W*_*in*_ *∈ ℝ*^*d*×1^ represents a shared projection matrix, *p*_*j*_ *∈ ℝ*^*d*×1^ represents a learnable positional encoding, and *σ* (*·*) denotes the sigmoid function. The learnable parameter *θ*_*j*_ generates a salience score *α*_*j*_ *∈* (0, 1), which acts as a soft filter. This mechanism allows the model to dynamically prioritize biologically relevant proteins while suppressing background noise specific to the disease context.

Knowledge-guided aggregation. We then aggregate these modulated protein embeddings into functional pathway representations using a prior knowledge mask *M ∈ {*0, 1*}*^*K,P*^ derived from the ConsensusPathDB^[15]^, where *K* = 333 denotes the number of biological pathways. If protein *j* is a component of pathway k, then *M*_*kj*_ = 1; otherwise *M*_*kj*_ = 0. The representation for the *k*-th pathway, denoted as *t*_*k*_, is computed by:

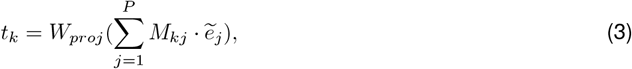

where *W*_*proj*_ *∈ ℝ*^*d*×*d*^ represents a linear transformation that refines the aggregated features. This formulation effectively transforms microscopic molecular signals into macroscopic functional tokens.

To ensure comprehensive coverage, proteins not annotated in the knowledge base (denoted as the set *U*) are processed via a supplementary parallel branch. Unlike the pathway tokens, these unannotated proteins *x*_*U*_ *∈ ℝ*^|*U*|^ are not encoded individually. Instead, they are projected via a global transformation to capture latent biological signals. Specifically, the concatenated vector of unannotated proteins is mapped through a dense layer and reshaped into a sequence of *A* auxiliary tokens 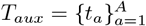 :

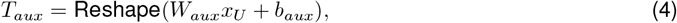

where *W*_*aux*_ *∈ ℝ* ^(*A·d*)*×*|*U*|^ projects the high-dimensional unannotated input into a latent feature space. The operator Reshape(*·*) transforms this flat vector from ℝ^*A·d*^ into a sequence structure ℝ^*A×d*^, effectively partitioning the latent features into *A* distinct tokens. These auxiliary tokens are then concatenated with the pathway tokens to form the complete sequence input for the Transformer.

##### 2.3.2.2. Modeling systemic interactions via Transformer

To capture the high-order systemic context, ProMeta processes the sequence of pathway tokens using a Transformer encoder^[16]^.

We construct the input sequence *Z*^(0)^ by concatenating a global classification token (*z*_*cls*_), the pathway tokens 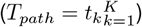, and the auxiliary tokens (*T*_*aux*_):

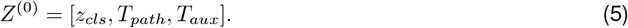

This sequence is processed through *L* layers of a standard Transformer. At each layer *l*, a multi-head self-attention (MSA) mechanism updates the representation of each pathway based on its interaction with all other pathways:

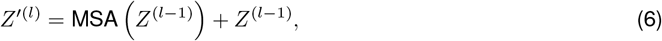

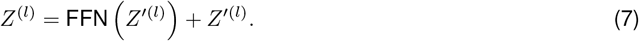

We also incorporate a global residual shortcut into the final representation to provide complementary feature signals, creating a dual-stream information flow. The resulting disease embedding *h*_*final*_ is the summation of the high-order pathway context and a linear projection of the raw proteomic profile:

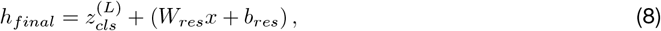

where *W*_*res*_ *∈ ℝ*^*d×P*^ projects the raw input to the latent dimension. The final prediction probability ŷ is obtained via a linear classifier:

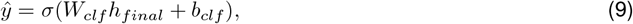

where *W*_*clf*_ *∈ ℝ*^1*×d*^ and *b*_*clf*_ *∈ ℝ* represent the classifier weights and bias, respectively.

#### 2.3.3. Meta-SGD optimization

##### 2.3.3.1 Task-adaptive bi-level optimization

To facilitate rapid generalization to novel conditions, we employ the Meta-SGD^[17]^ algorithm. This approach improves upon the classical Model-Agnostic Meta-Learning (MAML)^[18]^ framework by learning not only the model initialization but also the learning rate for each parameter. This capability enables more efficient and stable adaptation, a benefit that has been extensively demonstrated across diverse few-shot learning scenarios^[19]^. The training process is decoupled into two nested loops. Let *ϕ* denote the model parameters and *β* denote the learnable learning rate vector of the same dimension.

Inner loop (fast adaptation). For a sampled disease task 𝒯_*i*_ with a support set 𝒮_*i*_ = *{*(*x*_*s*_, *y*_*s*_)*}*, we update all model parameters *ϕ* on the support set *S*_*i*_ by performing five gradient steps with a learnable learning rate vector *β*:

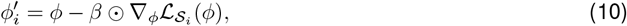

This allows the model to adjust the entire network to specific disease characteristics.

Outer loop (generalization). The adapted parameters 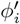 are evaluated on a query set 𝒬_*i*_ = {(*x*_*q*_, *y*_*q*_)}. The meta-objective minimizes the loss across a batch of tasks with respect to both the initialization *ϕ* and the learning rates *β*:

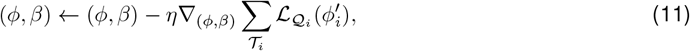

where *η* denotes the meta-learning rate.

##### 2.3.3.2. Objective function

Both the support loss 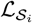 and query loss 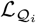 are computed using the same composite objective *L*, which enforces both the predictive accuracy and generalizability:

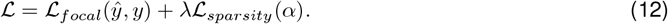

Here, ℒ_*focal*_ handles class imbalance by focusing on hard-to-classify examples^[20]^, defined as:

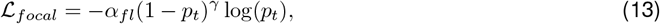

where *p*_*t*_ is the model’s estimated probability for the true class, *α*_*fl*_ is a balancing factor, and *γ* is the focusing parameter. The sparsity term 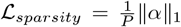 imposes a mild L1 penalty on the gating coefficients *α*, weighted by *λ*, to encourage the selection of salient biomarkers.

### 2.4. Experimental design and evaluation

#### 2.4.1. Meta-learning task construction

To evaluate the model’s capacity for strong generalization across both novel tasks and unseen individuals, we implemented a rigorous dataset splitting strategy. First, we partitioned the cohort into training (*P*_*train*_), validation (*P*_*valid*_), and testing (*P*_*test*_) sets using an 8:1:1 ratio. This strictly ensures that the support and query sets used during meta-testing are derived from individuals never observed during the meta-training phase.

We constructed the task distributions by allocating disease terms based on case availability within these population splits. Disease terms exhibiting *>* 50 and *<* 1000 positive cases within *P*_*test*_ were reserved for the testing task set 𝒯_*test*_. The remaining terms were allocated to validation (𝒯_*valid*_) and training (𝒯_*train*_) sets based on their prevalence in the validation and training cohorts, respectively. A threshold of 50 cases was selected to ensure sufficient sample sizes for conducting experiments across varying shot settings (up to 32-shot) while maintaining a robust query set for accurate performance evaluation.

To prevent information leakage arising from biological comorbidity, where related diseases share significant patient overlap, we employed a Jaccard-based pruning strategy. Let *S*_*t*_ and 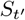 denote the sets of positive case identifiers for a training task *t* and a testing task *t*^*′*^. We explicitly removed any training task *t* exhibiting excessive overlap (*>* 75%) with any held-out testing task:

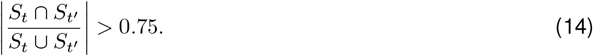

The final task distribution for prevalent analyses included 783 training, 23 validation, and 91 testing diseases; incident analyses included 972, 83, and 122 diseases, respectively. In contrast to selecting only a few specific diseases with scarce cases, this task construction method enables a more comprehensive and statistically robust evaluation.

During the testing phase on novel diseases (𝒯_*train*_ *∩ 𝒯*_*test*_ = ∅), the model utilizes the fixed global initialization *ϕ* and learning rates *β* learned during meta-training. For a given test disease, it performs Inner Loop adaptation on the provided support set 𝒮_*test*_ to derive task-specific weights 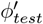 :

During the testing phase on novel diseases (𝒯_train_ *∩ 𝒯*_test_ = ∅), the model utilizes the meta-learned global initialization and learning rates *β*. Crucially, to ensure consistency with the training phase, we strictly adhere to the partial adaptation strategy. For a given test disease, the inner loop restricts updates to the adaptive parameters of classification head and the adaptive feature module (collectively denoted as *ϕ*_adapt_) using the support set *S*_test_, while the shared parameters *ϕ*_shared_ remain frozen to maintain universal biological representations. The task-specific weights 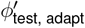 are derived as follows:

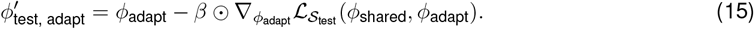

These adapted weights, combined with the frozen shared parameters, are then used to predict diagnoses for all samples in the query set 𝒬_test_.

#### 2.4.2. Model evaluation and benchmarking

To the best of our knowledge, no existing methods address disease modeling with extremely scarce cases using proteomics data. Consequently, we constructed several competitive baseline methods to systematically evaluate the meta-learning performance of ProMeta. We designed a suite of transfer learning-based strategies: (1) Multitask Transformer, which employs multi-task learning across all support sets in 𝒯_*test*_; (2) Supervised Transformer, utilizing supervised pre-training followed by fine-tuning on the support set of each disease in 𝒯_*test*_; (3) Self-supervised Transformer, utilizing self-supervised pre-training followed by fine-tuning; and (4) TaskSimilarity Transformer, where the model is first pre-trained on the most similar task in 𝒯_*train*_ before being fine-tuned on the corresponding target support set in 𝒯_*test*_. Additionally, we evaluated standard machine learning baselines, including Multi-Layer Perceptron (MLP), Random Forest, L1-regularized Logistic Regression (L1Logistic), and XGBoost^[21]^. Detailed implementations of these baselines are provided in the Supplementary Methods.

Binary classification performance was assessed using the Area Under the Receiver Operating Characteristic curve (AUROC) and the Area Under the Precision-Recall Curve (AUPRC). To ensure statistical robustness, performance estimates were derived from five independent runs with different random seeds for both data partitioning and model initialization. Crucially, identical training, validation, and testing splits were utilized across ProMeta and all baseline models to guarantee rigorous comparison.

We evaluated performance across 4-, 8-, 16-, and 32-shot scenarios. In each setting, the support set 𝒮_*test*_ was balanced (1:1 case-control ratio). For example, in the 4-shot setting, the support set contains 2 cases and 2 controls. To accurately evaluate generalization, the query set 𝒬_*test*_ was fixed at 128 samples with a reasonable class imbalance (approximately 25% cases and 75% controls). In instances where sufficient positive cases were unavailable to meet this ratio, additional controls were supplemented to maintain the total sample size of 128. Crucially, to ensure a fair performance comparison, the exact same query set was utilized across all varying support set sizes.

## Results

### 3.1. Disease diagnosis in few-shot scenarios

We first benchmarked the performance of ProMeta in diagnosing prevalent diseases. Across all experimental settings (ranging from 4-shot to 32-shot), ProMeta consistently and significantly outperformed all baseline methods, including both transfer learning and traditional machine learning approaches (Fig. 2). Even in the highly challenging 4-shot scenario (utilizing only 2 cases and 2 controls), ProMeta demonstrated robust generalization capabilities, achieving a mean AUROC of 0.693 and a mean AUPRC of 0.491 (Fig. 2a,b). In contrast, the best-performing baseline yielded a mean AUROC of only 0.556 (Supervised Transformer) and a mean AUPRC of 0.403 (MLP) (Fig. 2a,b). Subsequent experiments indicated a positive correlation between model performance and data availability: under the 32-shot setting, the mean AUROC and AUPRC of ProMeta climbed to 0.722 and 0.517, respectively.

**Fig. 2:**
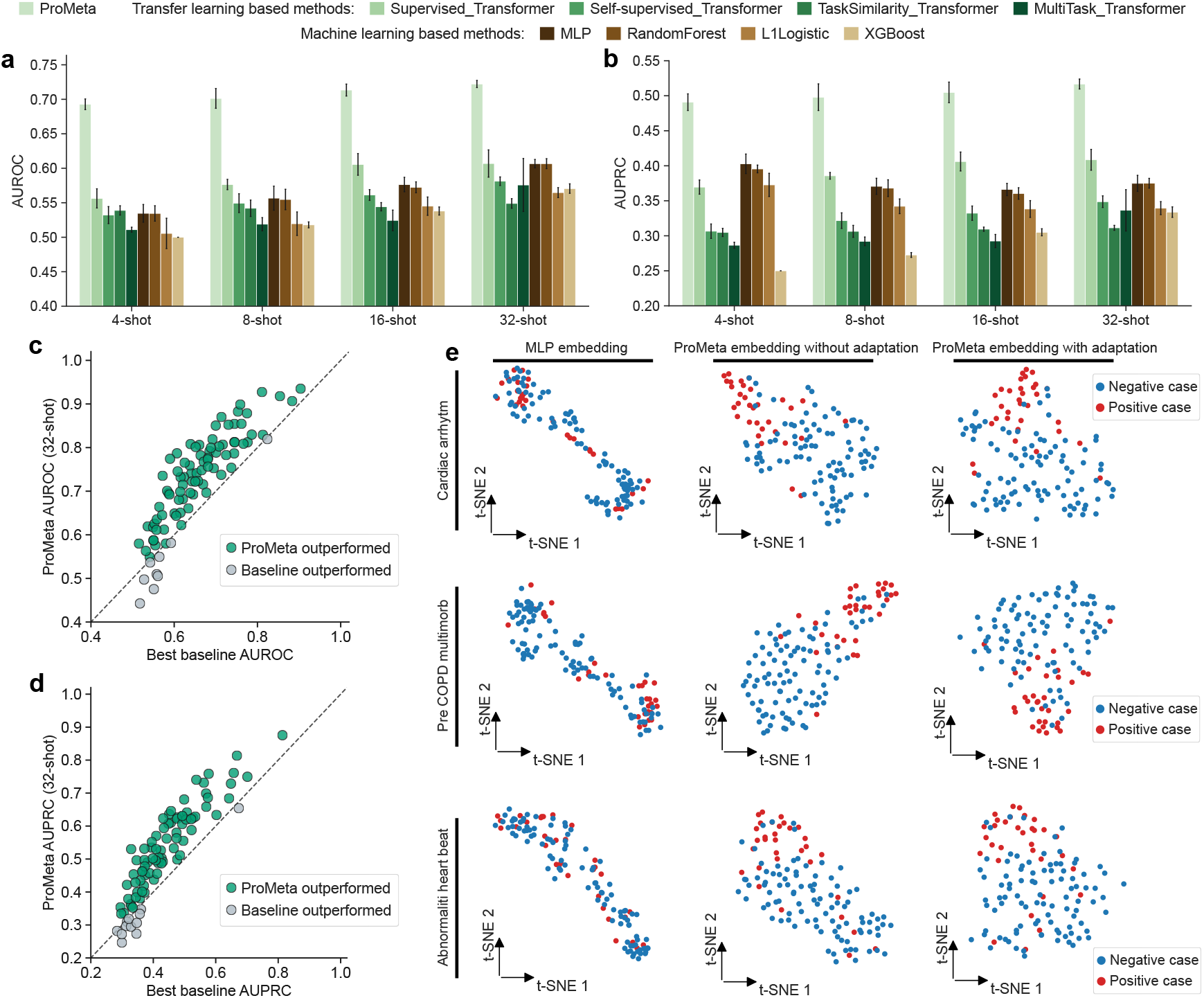
Performance evaluation for disease diagnosis in few-shot scenarios. (a-b) Overall (a) AUROC and (b) AUPRC scores comparing ProMeta against eight baseline methods across all testing diseases. The bar plot and error bars denote the mean and standard deviation, respectively, calculated across five independent experimental runs with different random seeds. (c-d) Head-to-head comparison of (c) AUROC and (d) AUPRC between ProMeta and the best-performing baseline method for each specific disease task. Each dot represents a unique disease endpoint. Red dots denote cases where ProMeta outperformed the best baseline method, while gray dots indicate cases where the best baseline outperformed ProMeta. The diagonal dashed line represents equal performance. (e) t-SNE visualization of sample embeddings generated by MLP (left), ProMeta prior to adaptation (middle), and ProMeta following adaptation (right) for selected disease classes. This visualization was performed on unseen testing tasks with a support set size of 32 (16 cases, 16 controls). AUROC, area under the receiver operating characteristic curve; AUPRC, area under the precision-recall curve; MLP, multilayer perceptron; t-SNE, t-distributed stochastic neighbor embedding.

At the individual disease level, ProMeta surpassed the respective best-performing baseline in 90.1% of diseases for AUROC and 86.8% for AUPRC (32-shot setting) (Fig. 2c,d). Interestingly, the proportion of tasks where ProMeta achieved superior performance increased with the support set size, demonstrating its superior modeling capacity (Supplementary Fig. 1a-d).

To investigate the source of ProMeta’s superiority, we visualized the latent embedding space. Results revealed that, unlike the MLP baseline which failed to distinguish samples, ProMeta effectively separated cases from controls even prior to task-specific adaptation (Fig. 2e). This confirms that the model’s global meta-initialization successfully captures biological priors generalizable to unseen diseases. Building on this foundation, adaptation with just a few shots further refined these representations, leading to substantial performance gains (Supplementary Fig. 1e).

Collectively, these results validate ProMeta as a robust framework for disease diagnosis, offering particular value for rare diseases where clinical data is inherently scarce.

### 3.2. Disease prediction in few-shot scenarios

The performance advantage of ProMeta extended seamlessly to the more challenging task of incident disease prediction (Fig. 3). Across the entire few-shot spectrum – from the data-scarce 4-shot to the 32-shot setting – ProMeta significantly outperformed both transfer learning and standard machine learning approaches. Quantitative metrics mirrored previous findings: starting with a mean AUROC of 0.697 and AUPRC of 0.464 in the 4-shot setting, performance scaled to 0.710 (AUROC) and 0.477 (AUPRC) in the 32-shot setting (Fig. 3a,b). These results underscore a strong positive correlation between support set size and predictive accuracy. Similarly, at the granular task level, ProMeta proved superior in the vast majority of endpoints, outperforming the best baseline in 91.0% (AUROC) and 82.0% (AUPRC) of tasks under the 32-shot setting (Fig. 3c,d; Supplementary Fig. 2a-d).

**Fig. 3:**
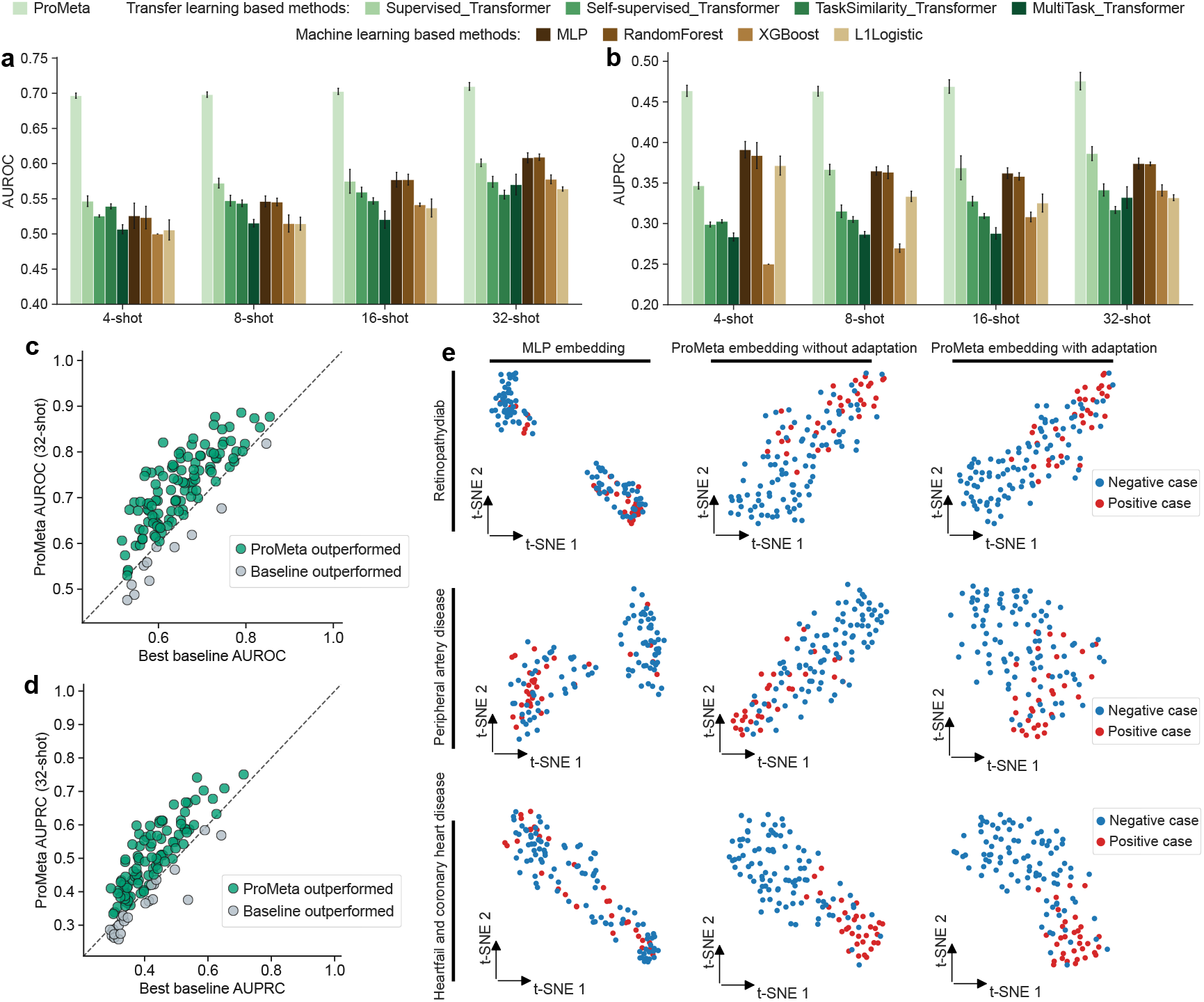
Performance evaluation for disease prediction in few-shot scenarios. (a-b) Overall (a) AUROC and (b) AUPRC scores comparing ProMeta against eight baseline methods across all testing diseases. The bar plot and error bars denote the mean and standard deviation, respectively, calculated across five independent experimental runs with different random seeds. (c-d) Head-to-head comparison of (c) AUROC and (d) AUPRC between ProMeta and the best-performing baseline method for each specific disease task. Each dot represents a unique disease endpoint. Red dots denote cases where ProMeta outperformed the best baseline method, while gray dots indicate cases where the best baseline outperformed ProMeta. The diagonal dashed line represents equal performance. (e) t-SNE visualization of sample embeddings generated by MLP (left), ProMeta prior to adaptation (middle), and ProMeta following adaptation (right) for selected disease classes. This visualization was performed on unseen testing tasks with a support set size of 32 (16 cases, 16 controls). AUROC, area under the receiver operating characteristic curve; AUPRC, area under the precision-recall curve; MLP, multilayer perceptron; t-SNE, t-distributed stochastic neighbor embedding.

Finally, visualization of the embedding space recapitulated the findings from the prevalent task. Unlike the entangled representations of the MLP baseline, ProMeta established clear separation between cases and controls prior to adaptation (Fig. 3e; Supplementary Fig. 2e). This confirms that the biological priors captured during meta-training are temporally robust, facilitating effective adaptation for future disease prediction.

### 3.3. Model interpretation

Beyond disease diagnosis and prediction, ProMeta facilitates the identification of disease-associated proteins and pathways. To achieve this, we introduced a gradient-based attribution method tailored for meta-learning (Supplementary Methods), which quantifies the specific contribution of each protein or pathway feature to a given disease phenotype.

For disease diagnosis tasks, ProMeta not only found a group of disease-associated proteins shared across different diseases, including FOLR3, GAST, and PAEP (Fig. 4a), but also revealed distinct clusters of proteins that corresponded to biologically related diseases (Supplementary Fig. 3). Furthermore, in the analysis of cardiovascular disease tasks, ProMeta assigned high importance scores to NPPB (Natriuretic Peptide B) (Fig. 4b), a well-established clinical gold-standard biomarker for heart failure^[22]^. Moreover, pathway-level analysis highlighted key pathogenic mechanisms, such as inflammation (e.g., the cytokine-cytokine receptor interaction pathway) and cell signaling dysregulation (e.g., PI3K-Akt and MAPK pathways) (Supplementary Fig. 4).

**Fig. 4:**
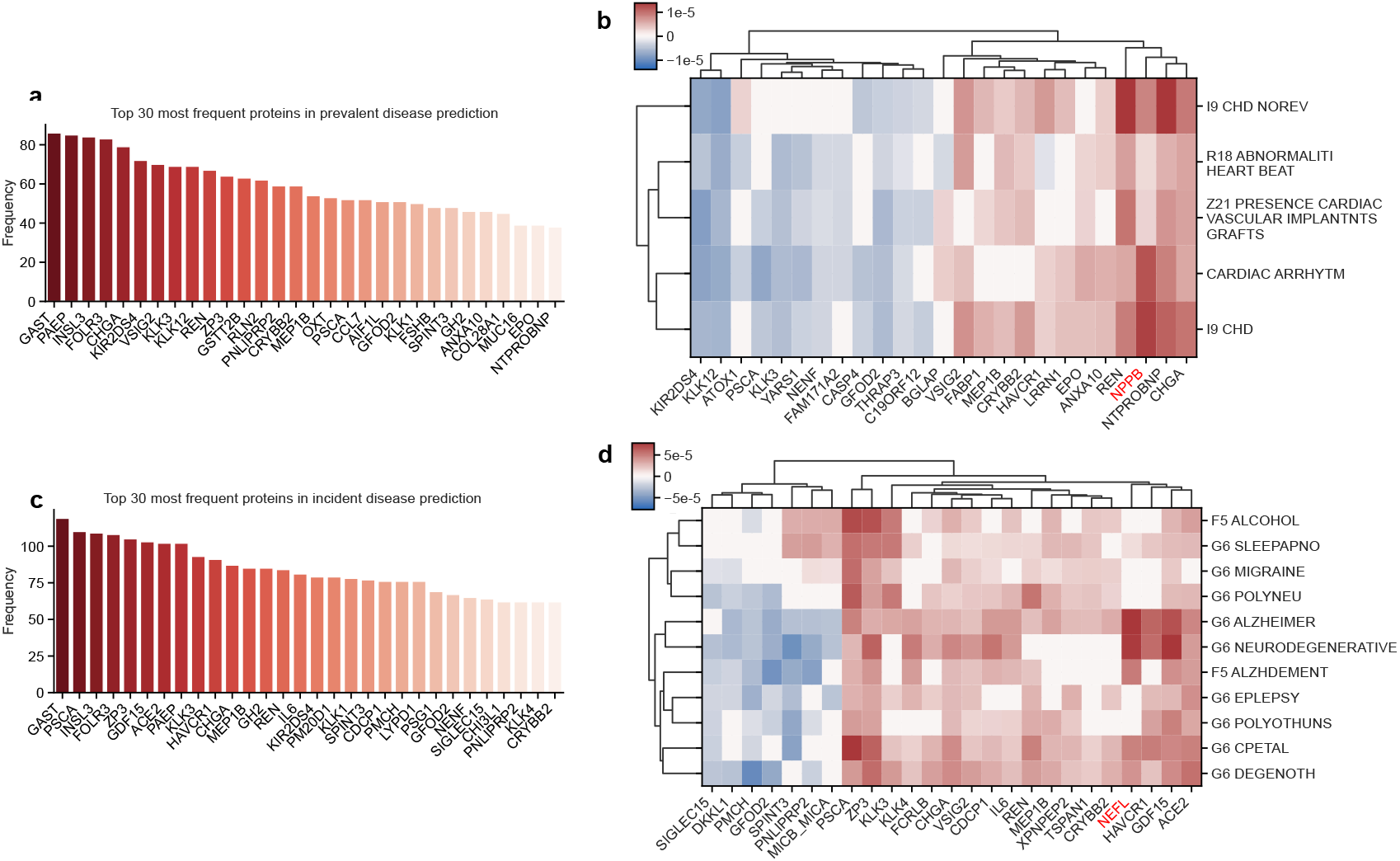
Biological interpretability of ProMeta. (a) Ranking of the top 30 most recurrent protein biomarkers across all prevalent diseases in 𝒯_*test*_. Frequency is defined as the number of unique disease tasks in which a protein appeared within the top-50 importance list. (b) Unsupervised hierarchical clustering of top-ranked protein biomarkers for major cardiometabolic diseases. Importance scores were derived from the ProMeta model trained on prevalent cases. (c) Ranking of the top 30 most recurrent protein biomarkers across all incident diseases in 𝒯_*test*_. Frequency is defined as the number of unique disease tasks in which a protein appeared within the top-50 importance list. (d) Unsupervised hierarchical clustering of top-ranked protein biomarkers for neurological and neurodegenerative disorders, with importance scores derived from the ProMeta model trained on incident cases. In both panels, importance scores are determined via ProMeta’s gradient-based attribution, where red indicates a positive association and blue indicates a negative association.

For disease prediction tasks, distinct from diagnostic tasks, ProMeta identified several prognostic regulators such as ACE2 and GDF15 ^[23]^ (Fig. 4c and Supplementary Fig. 5). This suggests that for future disease prediction, ProMeta prioritizes markers of physiological stress and homeostatic regulation. Crucially, when applied to predict future neurodegenerative risks, ProMeta identified NEFL (Neurofilament light chain) as a top predictor (Fig. 4d), consistent with its clinically established role as a biomarker for axonal damage and neuronal loss^[24]^. This protein-level finding is strongly corroborated by pathway-level analysis, which reveals a concurrent suppression of neuroprotective signaling (PI3K-Akt pathway) and dysfunction of GABAergic synapses (Supplementary Fig. 6). In summary, interpretability analysis highlights ProMeta’s robust ability to identify potential biomarkers for complex diseases.

### 3.4. Ablation study

In the preceding results, we demonstrated the effectiveness of our proposed method, where the meta-learning strategy showed significant superiority compared to alternative training paradigms (Fig. 2 and Fig. 3). To isolate the specific contribution of the ProMeta model architecture, we conducted ablation studies by replacing our backbone architecture with a standard Transformer (Transformer MAML) and an MLP (MLP MAML), while keeping the meta-learning strategy. Consistent with our initial findings, all meta-learning-based variants outperformed non-meta-learning baselines, further validating the efficacy of meta-learning for disease modeling under extreme data scarcity (Fig. 5). Crucially, ProMeta consistently outper-formed both Transformer MAML and MLP MAML across nearly all settings, confirming the added value of our proposed hierarchical dual-view modeling strategy (Fig. 5). Furthermore, we investigated the sensitivity of ProMeta to the number of auxiliary tokens used for unmapped proteins; the results indicate that the model’s predictive performance remains robust regardless of the token count (Supplementary Fig. 7).

**Fig. 5:**
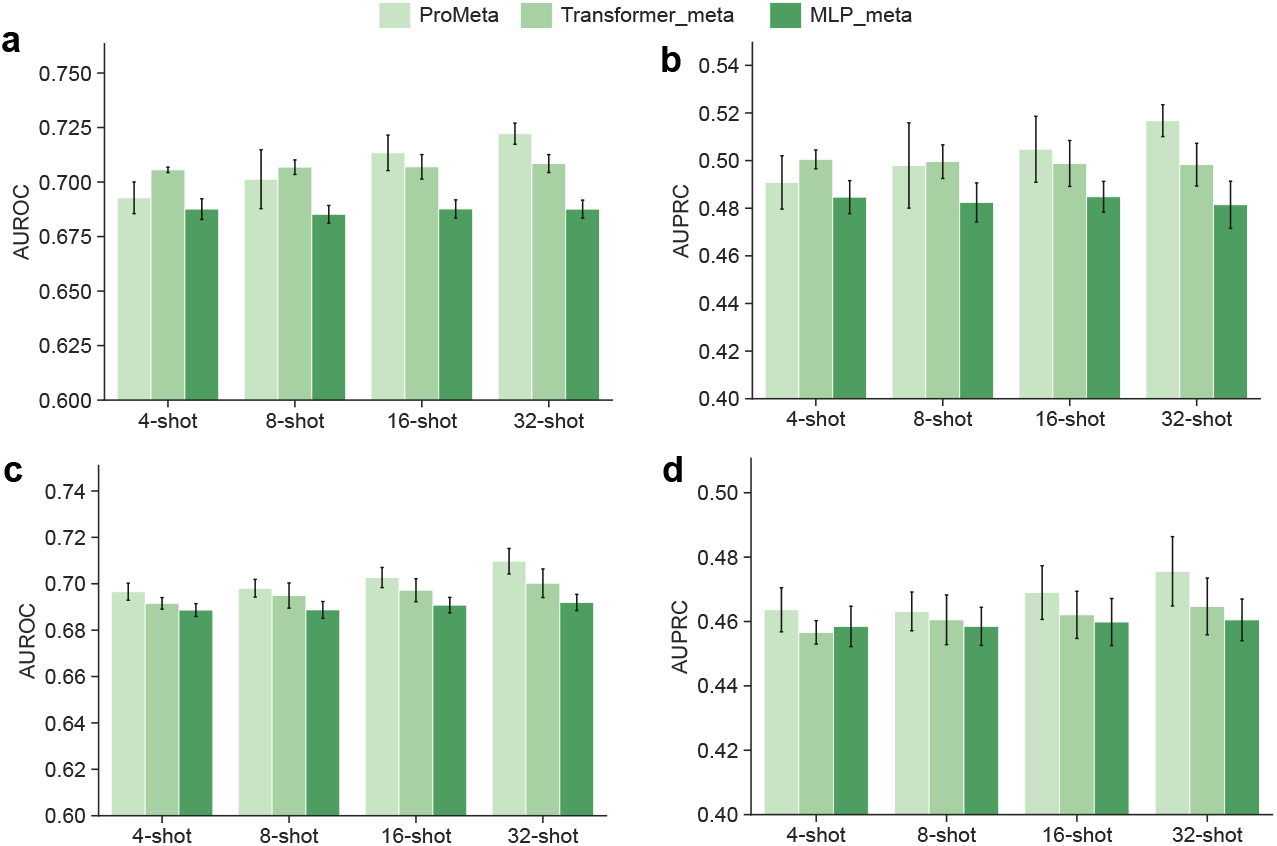
Ablation study of ProMeta. (a-b) Overall (a) AUROC and (b) AUPRC scores comparing ProMeta against the meta-learning baselines with MLP and Transformer backbones across all testing prevalent diseases. The bar plot and error bars denote the mean and standard deviation, respectively, calculated across five independent experimental runs with different random seeds. (c-d) Overall (c) AUROC and (d) AUPRC scores comparing ProMeta against the meta-learning baselines with MLP and Transformer backbones across all testing incident diseases.

## Discussion

In this study, we introduced ProMeta, a prior knowledge-guided meta-learning framework that addresses a fundamental bottleneck in proteomic precision medicine: the failure of current deep learning models to generalize to conditions with scarce data. By synergizing a pathway-aware transformer architecture with bi-level optimization, ProMeta successfully extracts transferable biological priors from biobank-scale datasets, enabling robust disease modeling even in extreme few-shot scenarios. Our extensive benchmarking across 91 prevalent and 122 incident disease conditions demonstrates that ProMeta consistently outperforms the state-of-the-art transfer learning and traditional machine learning baselines. The model’s ability to achieve high diagnostic and prognostic accuracy with as few as four patient samples validates the efficacy of the “learning-to-learn” paradigm in proteomic analysis. Mechanistically, we show that ProMeta does not merely memorize task-specific patterns; rather, it learns a universal latent initialization that almost linearly separates physiological states prior to model adaptation. Furthermore, ProMeta offers inherent interpretability, identifying key proteins and pathways that align with established pathophysiology. In summary, ProMeta represents a pivotal step toward a personalized proteomic analysis framework.

## Data and code availability

All proteomic, phenotypic, and EHR data used in this study are available from UKB upon application (https://www.ukbiobank.ac.uk).

The source code of ProMeta is available at GitHub (https://github.com/lihan97/ProMeta).

## Acknowledgments

This work was supported by the National Key R&D Program of China (2024YFC3407800 to H.L.) and Natural Science Foundation of Tianjin (25JCQNJC01180 to H.L.)

## Competing interests

No competing interest is declared.

## Author contributions

H.L. and S.Z. conceived the concept and designed the study. H.L., H.G. and L.H. developed ProMeta and performed data analysis. H.L., H.G., L.H., Y.L., Z.Z., P.G., J.C.-K., Y.M., J.Z. and S.Z. are responsible for data interpretation. S.Z., J.Z., Y.M. and H.L. supervised the project. H.L., H.G., L.H. and S.Z. prepared the manuscript with assistance from all other authors.

## Notes

### Competing Interest Statement

The authors have declared no competing interest.

